# NetAn: A Python Toolbox for Network-based Gene Annotation Enrichment

**DOI:** 10.1101/2023.09.05.556339

**Authors:** Eduard Ghemes, Zelmina Lubovac-Pilav, Rasmus Magnusson

## Abstract

**Background:** Gene annotation enrichment analysis is the gold standard for studying the biological context of a set of genes, but available tools often overlook important network properties of the underlying gene regulatory system.

**Results:** We present the NETwork ANnotation enrichment package, NetAn, built in Python, which augments annotation analysis with approaches such as inference of closely related genes to include local neighbors in the analysis, the extraction of separate network sub-clusters, and the following over-representation analyses based on network clustering. By using NetAn, we demonstrate how these approaches enhance the identification of relevant annotations in human gene sets. In a specific case study on Multiple Sclerosis (MS), NetAn’s approach of incorporating neighboring genes through network-based expansion demonstrates a distinct advantage in identifying immune-related genes critical to MS pathology. Furthermore, we demonstrate the ability of NetAn to stratify MS annotations to also identify relevant neuron-related enrichments. Lastly, we compare NetAn to alternative network-based approaches, and find it to have greater specificity compared to broader approaches like NET-GE.

**Conclusions:** We present NetAn, a novel network-based approach that can stratify annotation enrichment analyses by integrating gene interactions and network topology, thereby strengthening biological signals through the inclusion of associated genes. This approach allows for enhanced identification of disease-relevant annotations, as demonstrated in the MS case study.

## Introduction

Gene annotation enrichment analysis is the cornerstone approach for establishing the association of biological functionality with a set of investigated genes. This allows users to pinpoint which biological processes, pathways, or molecular functions are overrepresented in a set of genes, helping to uncover mechanisms that might contribute to, as an example, disease progression. The basic idea of gene enrichment analysis is to systematically identify biological terms that are relevant towithin the provided gene set. While several tools for identifying annotations enrichment on gene sets have been developed (Kanehisa et al., 2016; Aleksander et al., 2023; Magnusson & Lubovac-Pilav, 2021), most of them encounter challenges related to at least two key factors.

Firstly, if the studied gene set is associated with multiple biological processes, the signal will effectively be obscured in each individual statistical test, increasing the risk of false negative identifications. Secondly, genes form a complex network; yet the topology of this network is frequently overlooked in gene set annotation enrichment analyses. Many complex diseases, especially those in which multiple processes are dysregulated simultaneously, such as multiple sclerosis (MS), could greatly benefit from a more comprehensive enrichment analysis.

MS is a chronic autoimmune disorder affecting the central nervous system (CNS), leading to different complications involving chronic inflammation, demyelination and finally neuronal damage (Attfield et al., 2022). By performing enrichment analysis on MS gene sets, valuable insights into disease’s pathogenesis can be uncovered. Since MS is a notoriously complex disease, by performing a simple enrichment analysis, some biological terms from less represented groups will arguably be overshadowed in the final results. As such, annotation enrichment analysis methods that account for topologically related genes are needed to both correctly identify and disentangle related processes.

To address these limitations, we developed the Network Annotation Enrichment (NetAn) package, a Python toolkit designed to integrate network perspectives into gene annotation enrichment analysis. NetAn improves upon traditional annotation tools by considering the relationships between genes and their place in biological networks, offering a more accurate identification of enriched terms in complex diseases. This type of analysis can handle the complexity of gene interactions, ensuring that even less represented genes and pathways are not overlooked.

We validated NetAn’s performance using MS-related gene sets, and highlight the network-based contributions of key genes involved in MS. Furthermore, we compared the results to existing tools such as NET-GE (Bovo et al., 2016) and netGO (Kim et al., 2020), finding that NetAn provides more specific and biologically plausible enrichment compared to broader approaches, such as NET-GE.

## Implementation

The core of NetAn regards how gene annotation enrichment analyses can be augmented by *a priori* defined biological networks. While NetAn provides built-in support for GO annotations (Aleksander et al., 2023), KEGG pathways (Kanehisa & Goto, 2000) and a processed STRINGdb network (Szklarczyk et al., 2015), users can also provide custom annotation sets or networks for customized usage. An overview of the tool can be seen in **Fig. 1**.

**Figure 1:**
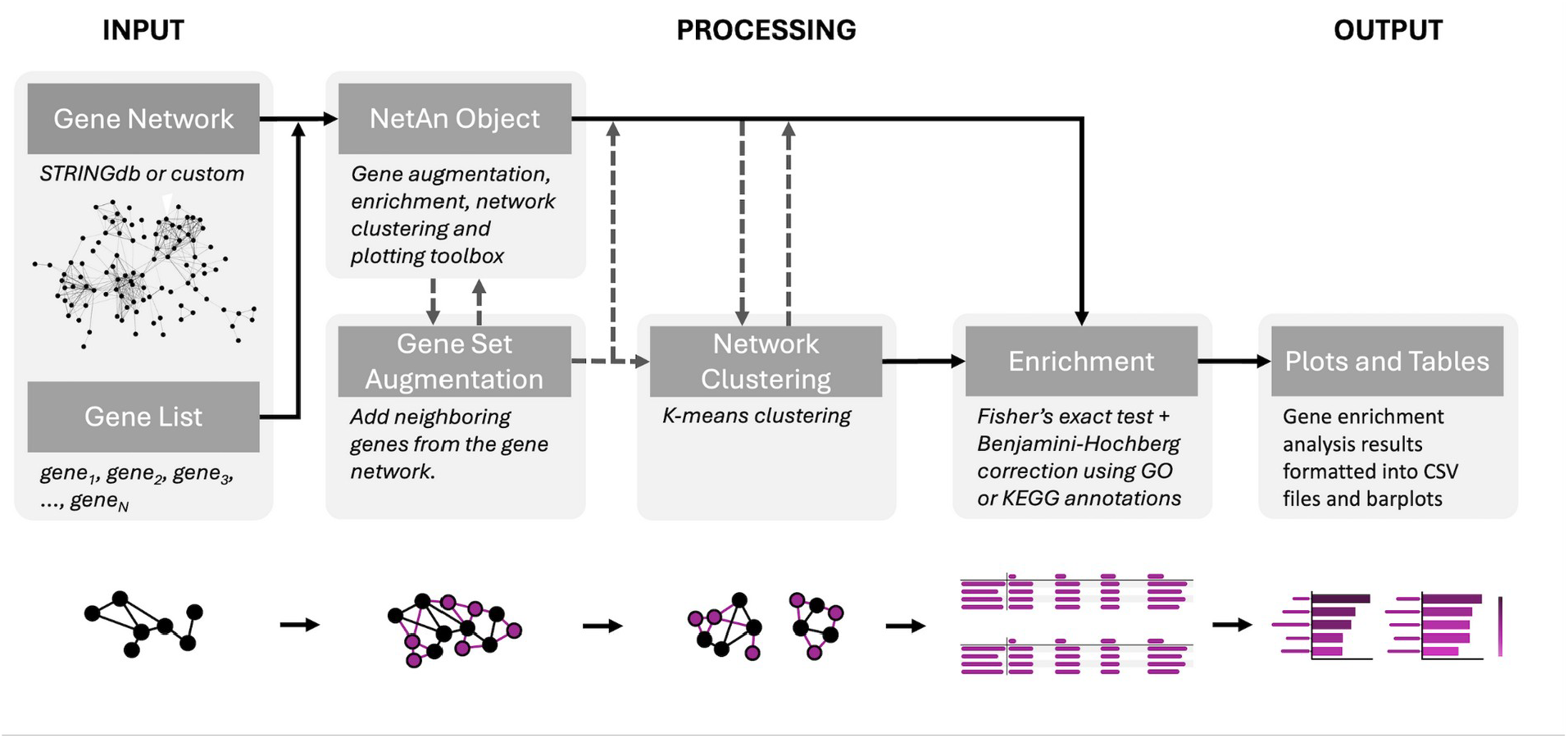
Overview of NetAn functionalities. Aa a first step in the workflow, a gene list and a network, such as the built-in STRINGdb network are provided as input. Optionally, gene set augmentation adds neighboring genes, and network clustering groups genes based on topological characteristics. This is followed by enrichment analysis using Fisher’s exact test with Benjamini-Hochberg correction based on GO, KEGG or custom annotations. The outputs are presented as plots and tables.

### Including topologically close genes in the annotation enrichment analysis

One of the main functionalities of NetAn is its ability to expand the set of studied genes for enrichment analysis based on topological proximity. Through this expansion, NetAn can add statistical power to a small initial set of genes by calculating the sum of edge weights for all connected nodes and then normalizes them against a STRINGdb network-derived background. NetAn next appends the top M*N associated genes, where M is the number of input genes with a STRINGdb ID and N is a factor of M (set by the user, default N=2.0).

### Identifying and analyzing topological gene cluster enrichments individually

While the use of network topology in annotation enrichment analyses has been explored before (Glaab et al., 2012; Bovo et al., 2020; Cousins et al., 2023), few approaches have used networks to disentangle complex cellular mechanisms in enrichment analyses. If the studied gene set involves genes from two or more moderately independent processes, the signal is easily obscured, and false negative identifications will be more likely. An optimal approach would instead divide the set of genes into distinct clusters based on their roles in different processes. NetAn accomplishes this by using network clustering techniques to identify clear divisions among the gene sets.

To this end, NetAn first constructs the adjacency matrix of the input gene set from the loaded network and then applies K-means clustering to group the network into a predetermined number of clusters, which is useful when dealing with complex gene networks encompassing several processes. Once the genes are grouped into these clusters, NetAn tests the observed versus expected frequency of gene associations within each set by performing a Fisher’s exact test-based annotation enrichment analysis on each cluster. Next, either the non-clustered (**Fig. 2A-B**) or clustered (**Fig. 2C**) results can be plotted using the in-build plotting function.

**Figure 2:**
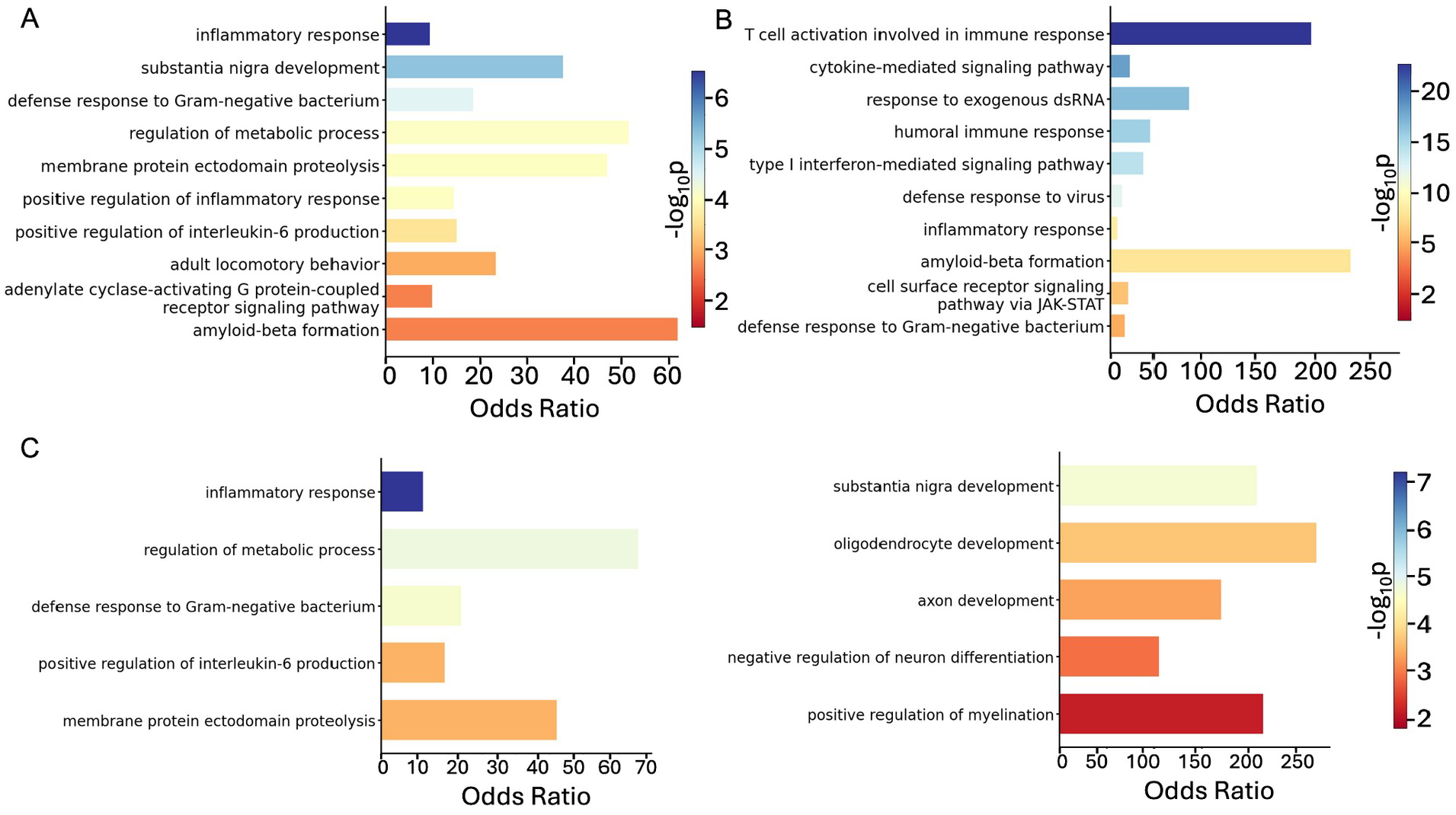
Enrichment analysis of original (A), expanded (B) and clustered (C) annotation enrichment analysis. The color gradient represents statistical significance, with bluer colors indicating a lower p-value. **(A)** The original gene set shows a predominant enrichment of inflammatory response and cytokine activity-related terms. **(B)** Expanding the analysis, using NetAn’s gene set expansion functionality to incorporate neighboring genes from the STRINGdb network reveals a more pronounced immune-related list of significantly enriched terms. **(C)** Shifting the focus to the clustered analysis, we can see clear separation between the groups of enriched terms. The bottom right panel focuses on neurobiologically-related processes, some of which were overshadowed in the non-clustered analysis. The bottom left panel highlights processes more similar to the ones that predominate the non-clustered analysis. This suggests that genes closely linked to the original set are predominantly involved in immune pathways, reinforcing the immune system’s relevance in MS.

### Measuring cluster differences by network average distances

To analyze the independence between identified cluster subsets, NetAn includes a functionality that computes the average shortest path length between all gene pairs between clusters. This average shortest path is compared to the average path length of the entire loaded network. The process can be broken down in three steps. First, it generates random positions and samples genes from each list. Second, it selects genes at these random positions and iterates over them to calculate distances. Third, it returns an array of the shortest paths between the genes. This comparison helps reveal the extent of connectivity between gene clusters, indicating potential biological relationships between seemingly distinct processes. The significance of the difference in these path lengths is tested using a non-parametric Mann-Whitney U test. For efficiency, if the number of pairs between the clusters is too large to calculate directly, NetAn uses the Monte Carlo method to randomly sample pairs in batches, calculating the averages until the mean stabilizes, providing a balance between accuracy and efficiency.

### Exporting and Visualization

NetAn provides a functionality to export results in the form of a csv-text file including annotations, odds ratios (ORs), nominal p-values, and multiple testing-corrected p-values. The package also offers a built-in plotting tool, with formatting options. In summary, a user of NetAn can go from a list of identified genes to a plot of sets of functionally independent gene sets with an annotation-based biological context, all in <5 lines of Python code.

## Results and discussion

### A Case Study of NetAn Applied to Multiple Sclerosis reveals distinct annotation sub-clusters

To demonstrate NetAn’s applicability to complex diseases, we applied it to genes associated with multiple sclerosis (MS). Specifically, we downloaded the 111 MS-associated genes from the *Diseases* database (Pletscher-Frankild et al., 2015). By performing an annotation enrichment analysis using NetAn, we found a strong prevalence of immunologically related annotations among the top results **(Fig. 2A)**. Genes associated with the inflammatory response and cytokine activity, which are characteristic traits of MS (Attfield et al., 2022), exhibit the strongest enrichment, as expected. Genes associated with myelin sheath and neurodevelopmental processes were also enriched, including terms like oligodendrocyte development and axon development **(Fig. 2C)**. Additionally, the other immune-related terms might underscore the involvement of immune-specific molecules, such as interleukins (e.g., IL-17, IL-23), key regulators of Th17 cell activity implicated in MS pathogenesis (Milovanovic et al., 2020; Li et al., 2016; Hiltensperger & Korn, 2018).

### Clustering

Applying the clustering functionality in NetAn on the MS-associated genes, revealed a sub-cluster consisting mainly of neurobiologically related annotation terms, while the first cluster remained focused on immune-related terms. The gene set within this new cluster yielded 13 significant terms (padj <0.05, Benjamini-Hochberg-adjusted), 11 of which were also significant in the non-clustered analysis. However, their rankings were significantly lower in the non-clustered analysis, with a median position of 34 compared to top positions observed in the clustered analysis. This might lead to potential oversight without network stratification of the gene set, greatly benefiting more complex diseases.

### Gene set expansion

Lastly, NetAn’s gene set expansion functionality was employed to augment the MS-related gene set. By doubling the initial 111 MS genes with topologically close genes in the network, the analysis yielded 26 new significant annotations (padj < 0.05, Benjamini-Hochberg-adjusted). In the expanded gene set, it is still evident that cytokines play a big role in MS **(Fig. 2B)**. Most of the enriched terms, such as cytokine-mediated signaling, are relevant to MS (Mezghiche et al., 2024; Milovanovic et al., 2020; Navarro-Compán et al., 2023; Bergamaschi et al., 2010; Yeo et al., 2007).

The full details about the genes associated with the annotation terms in each test can be found in the **Supplementary Materials**.

### Performance evaluation of NetAn

While gene network topologies are, arguably, often overlooked in annotation enrichment analyses, we sought to compare NetAn to other tools like NET-GE (Bovo et al., 2016), EnrichNet (Glaab et al., 2012) and netGO (Kim et al., 2020). NET-GE, EnrichNet and netGO enhances gene enrichment analysis by integrating protein-protein interaction networks The analysis comparing the genes that are the most associated with ontology terms between NetAn, EnrichNet and NET-GE revealed key differences in the most enriched terms and the genes associated with the said terms. Nonetheless, a few genes, such as *APP* (Amyloid Beta Precursor Protein), PSEN1 (Presenilin 1), and SNCA (Alpha-synuclein) were among the top genes associated with ontology terms in all the tested tools.

Among the tested tools, NET-GE results appear to diverge the most from those produced by NetAn, as measured by the correlation of the ranked enrichment terms in common for both NetAn and NET-GE (Spearman rho = -0.06). In contrast, EnrichNet identifies a set of annotations that are more closely aligned with NetAn (Spearman rho = 0.45), both in terms of annotation overlap and relative ranking **(Fig. 3)**. NetAn identified 39 significant biological processes out of a total of 1576 associations (∼2.7% significant terms). EnrichNet returns 1816 annotation terms, of which 55 are significant (∼3% significant terms, similar to NetAn). Meanwhile, NET-GE, in contrast, reports only significant annotations, yielding a total of 342 terms. This suggests that NetAn may provide a more balanced annotation profile than NET-GE, capturing relevant terms without excessive noise or loss of information. The convergence between NetAn and EnrichNet suggests that NetAn’s results are not only biologically meaningful but also reproducible, adding weight to the reliability of its findings.

**Figure 3:**
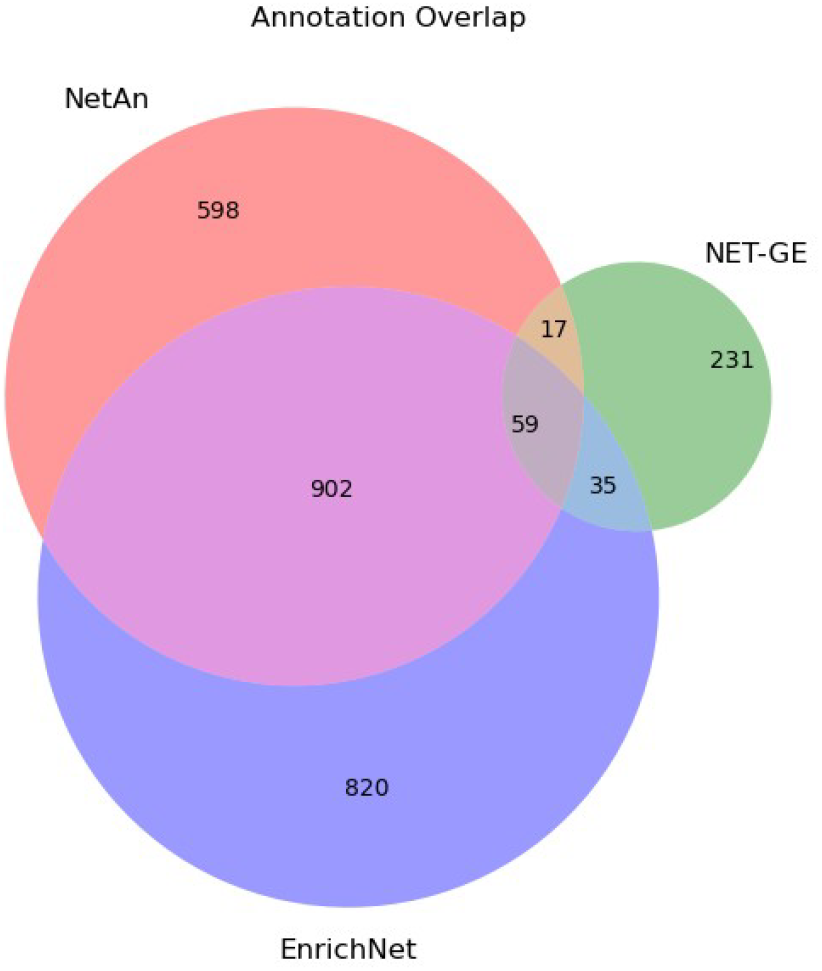
Venn diagram illustrating the overlap between all the identified annotations. Venn diagram illustrating the overlap between all the identified annotations. The highest overlap is seen between EnrichNet and NetAn results (Jaccard index of 0.4). In contrast, NET-GE identified a very small number of terms, with very few in common with the other tools (Jaccard index of 0.04 and 0.05, respectively).

Given that the gene set used in this example is associated with MS, the ranking of terms provided by NetAn appears to more accurately reflect the disease’s biological relevance. For instance, terms such as *“oligodendrocyte development”* rank much higher in NetAn compared to the other tools. Other terms, such as *“substantia nigra development”*, are entirely absent from the results of the other tools. NetAn, however, prioritizes processes more directly involved in MS pathology, thereby reducing the risk of introducing irrelevant annotations.

To more thoroughly compare NetAn’s performance with other network-based annotation enrichment tools, we also used the example dataset provided with the netGO tool (Kim et al., 2020). This dataset comprises nearly 330,000 gene-annotation pairs and includes 30 genes associated with BRCA breast cancer.

It is noteworthy that netGO takes 5-25 minutes for this application, while NetAn completes the task in seconds. Moreover, netGO identified 9 annotations after multiple testing corrections, all of which were associated with cancers in various tissues, including breast, colon, and lungs. In parallel, the standard application of NetAn identified two annotations, both of which were related to breast cancer and were also identified by netGO.

### Network augmentation increases statistical power

The next test examined NetAn’s ability to infer annotations from a broader gene set. We used the full Diseases database (Pletscher-Frankild et al., 2015), containing 477,770 gene-disease associations related to 5,776 distinct diseases, was considered. From this dataset, 100 diseases with more than 10 but fewer than 100 associated genes were randomly selected. Subsequently, NetAn was applied to predict annotations based on two scenarios: using the associated genes alone, and using the associated genes along with their local neighbors within the STRINGdb network (Szklarczyk et al., 2015), as integrated into NetAn.

Each analysis was typically executed within less than 60 seconds. This process was performed for various gene fold expansions, i.e. the fraction of the added genes in relation to the size of the original set, ranging from 0.5 to 2.5 in intervals of 0.5. To illustrate, a fold expansion of 0.5 using an initial set of 10 genes would yield an additional 5 genes. The results indicated that NetAn consistently augmented the number of significant annotations (padj <0.05, Benjamini-Hochberg corrected). For the multiple fold expansion inputs between 0.5 and 2.5, the number of significant annotations increased 1.3, 1.5, 2.7, 2.2, and 3.5-fold, respectively.

Importantly, annotations that were exclusively significant after gene expansion were near significance in the annotation enrichment analysis based on the non-expanded gene set. Median nominal p-values for these annotations were all significant, with median odds ratios (ORs) of 18.0, 16.1, 13.8, 11.1 and 11.7, respectively **(Fig. 4)**.

**Figure 4:**
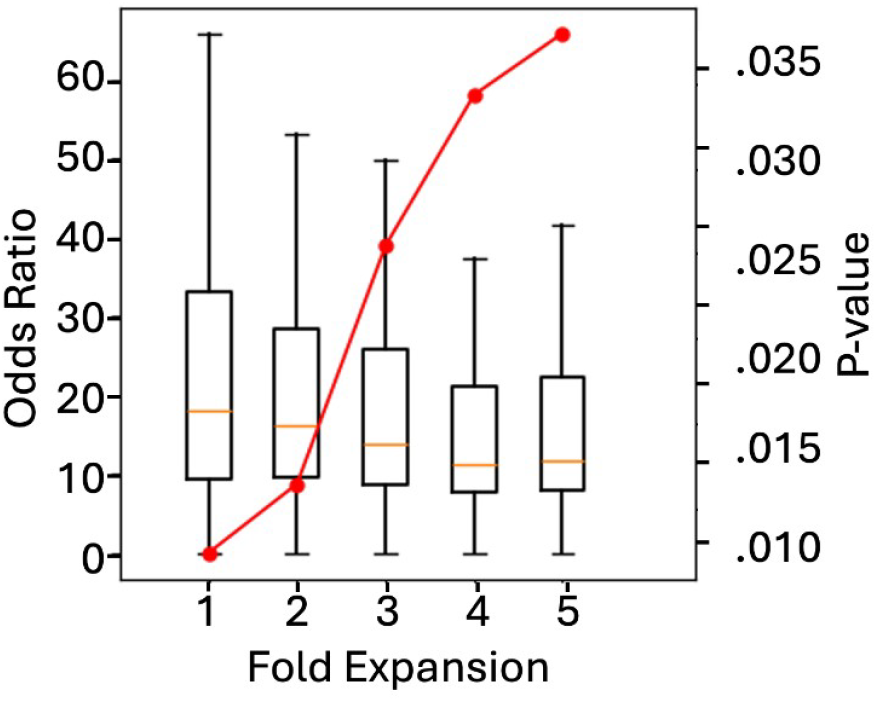
Multiple gene set expansions, with the associated p-values and odds ratios. By expanding the gene set tested in the ORA using NetAn, there was a clear increase in the number of significant annotations. The boxplots represent the distribution of ORs for each level of gene set expansion, while the red connected dots indicate the median p-values for the annotations at each fold expansion level. As the number of genes included in the ORA increases, the median odds ratios tend to decrease, and the median p-values tend to increase, although they remain significant.

## Conclusions

NetAn provides a robust and efficient framework for network-based gene annotation enrichment analysis, addressing key limitations of traditional methods. By integrating network topology and enabling functionalities such as gene set expansion and cluster-specific analysis, NetAn enhances the identification of biologically relevant annotations, particularly for complex diseases like multiple sclerosis. Its flexibility, speed, and specificity make it a valuable tool for researchers.

## Abbreviations

NetAn: NETwork ANnotation enrichment package
MS: Multiple Sclerosis
CNS: Central Nervous System
GO: Gene Ontology
KEGG: Kyoto Encyclopedia of Genes and Genomes
OR: Odds Ratio

## Declarations

## Ethics approval and consent to participate

Not applicable

## Consent for publication

Not applicable

## Availability of data and materials

NetAn was developed as a Python 3 package, and is available under a GNU General Public License V3. The package, along with a Jupyter Notebook tutorial, is freely available at https://github.com/Eduard-Ghemes/NetAn. NetAn is installed locally using the Python package management system pip, to allow for a simple and fast install.

Project name: NetAn

Project home page: https://github.com/Eduard-Ghemes/NetAn

Operating system(s): Platform independent

Programming language: Python 3, Jupyter

Other requirements: None

License: GNU General Public License V3

Any restrictions to use by non-academics: None

Data analyzed as part of this study: http://diseases.jensenlab.org/

## Author contributions

R. Magnusson – conceptualization, study design, package development, manuscript writing

E. Ghemes – package development, manuscript writing

Z. Lubovac-Pilav – conceptualization and study design

